# A structure-based model for the electrostatic interaction of the N-terminus of protein tau with the fibril core of Alzheimer’s Disease filaments

**DOI:** 10.1101/484279

**Authors:** David R. Boyer, David S. Eisenberg

## Abstract

Although portions of tau protein within the microtubule binding region have been shown to form the ordered core of tau filaments, the structural details of how other regions of tau participate in filament formation are so far unknown. In an attempt to understand how the N-terminus of tau may interact with fibril core, we crystallized and determined the structure of the N-terminal segment _5_RQEFEV_10_ of tau. Several lines of evidence have shown the importance of this segment for fibril formation. The crystal structure reveals an out-of-register Class 5 steric zipper with a wet and a dry interface. To examine the possible interaction of _5_RQEFEV_10_ with the tau fibril core, we modeled the binding of the wet interface of the _5_RQEFEV_10_ structure with the _313_VDLSKVTSKC_322_ region of the Alzheimer’s Disease tau filament structures. This model is consistent with, and helps to explain previous findings on the possible interaction of these two segments, distant in sequence. In addition, we discuss the possible conservation of this interaction across multiple polymorphs of tau.

## Introduction

The aggregation of tau into amyloid fibrils is associated with some 25 neurological diseases, collectively termed tauopathies. Although scientists have for decades associated fibrous tau aggregates with disease for decades, the molecular events driving aggregation of tau into amyloid fibrils remain unknown. It is generally thought that tau remains in three pools in the cell: attached to microtubules to promote their stability(1,2), bound to molecular chaperones to protect nucleating sequences of tau from enabling aggregation(3), or in a fibrous state where each fiber contains hundreds to many thousands of tau molecules(4–6). Under what conditions the fibril state begins to dominate is unclear.

Previous studies have shown that soluble, monomeric tau largely lacks a defined 3-dimensional shape(7); however, other studies posit that tau adopts a “paper clip” conformation in solution(8) or a seed-competent conformation where amyloid nucleating sequences are exposed and able to seed fibril formation(9). In addition, the binding of different tau constructs to microtubules has been visualized by cryo-EM(2). Despite these findings, information on the structure of soluble, monomeric form of tau is limited due to its largely disordered nature; therefore, most structural studies have focused on the aggregated state of tau(10–14). Our laboratory first focused on the segments of tau shown to be essential for *in vitro* aggregation, the primary nucleating sequences VQIINK and VQIVYK, located at the beginning of tau microtubule binding repeats 2 and 3, respectively(15). The crystal structures of these segments revealed classical “steric zipper” structural features(10,11). Mutations to these segments inhibit full-length tau aggregation, and we have shown that inhibitors designed to “cap” the crystal structures of VQIINK and VQIVYK segments also inhibit full-length tau aggregation, further demonstrating the importance of these segments(11,16).

Recently, cryo-EM studies of extracted tau filaments from Alzheimer’s Disease and Pick’s Disease patients have revealed several tau fibril polymorphs in near-atomic detail(12–14). In all of these structures, residues 306-378 spanning the length of Repeats 3 and 4 plus an additional six residues to the C-terminus of Repeat 4, are ordered in the fibril core, and in Pick’s Disease, Repeat 1 residues 254-274 are also ordered(14). Although these landmark discoveries help illuminate the fold adopted by the microtubule binding region of tau, it is still unknown to what degree other parts of tau participate in the aggregation process.

In the AD fibril structures, there is additional density consistently seen near residues K317 and K321 that may indicate another region of tau is interacting with the fibril core(12,13). Fitzpatrick *et al.* hypothesize that this extra density belongs to the residues _7_EFE_9_, an N-terminal sequence of tau that is part of the Alz50/MC-1 antibody binding epitope(12,17). To better understand the potential interaction of the N-terminus and the AD fibril core, we sought to determine the structure of this N-terminal segment.

## Results

We first searched for segments containing _7_EFE_9_ that are likely to crystallize. Although no segment containing _7_EFE_9_ scored well on the structure-based ZipperDB server(18), the ability to form fibrils from segment _5_RQEFEV_10_ was previously predicted by a sequence-based method and demonstrated biochemically(19). Therefore, we crystallized and determined the structure of the hexameric segment _5_RQEFEV_10_ (Figure 1 A-C).

**Figure 1:**
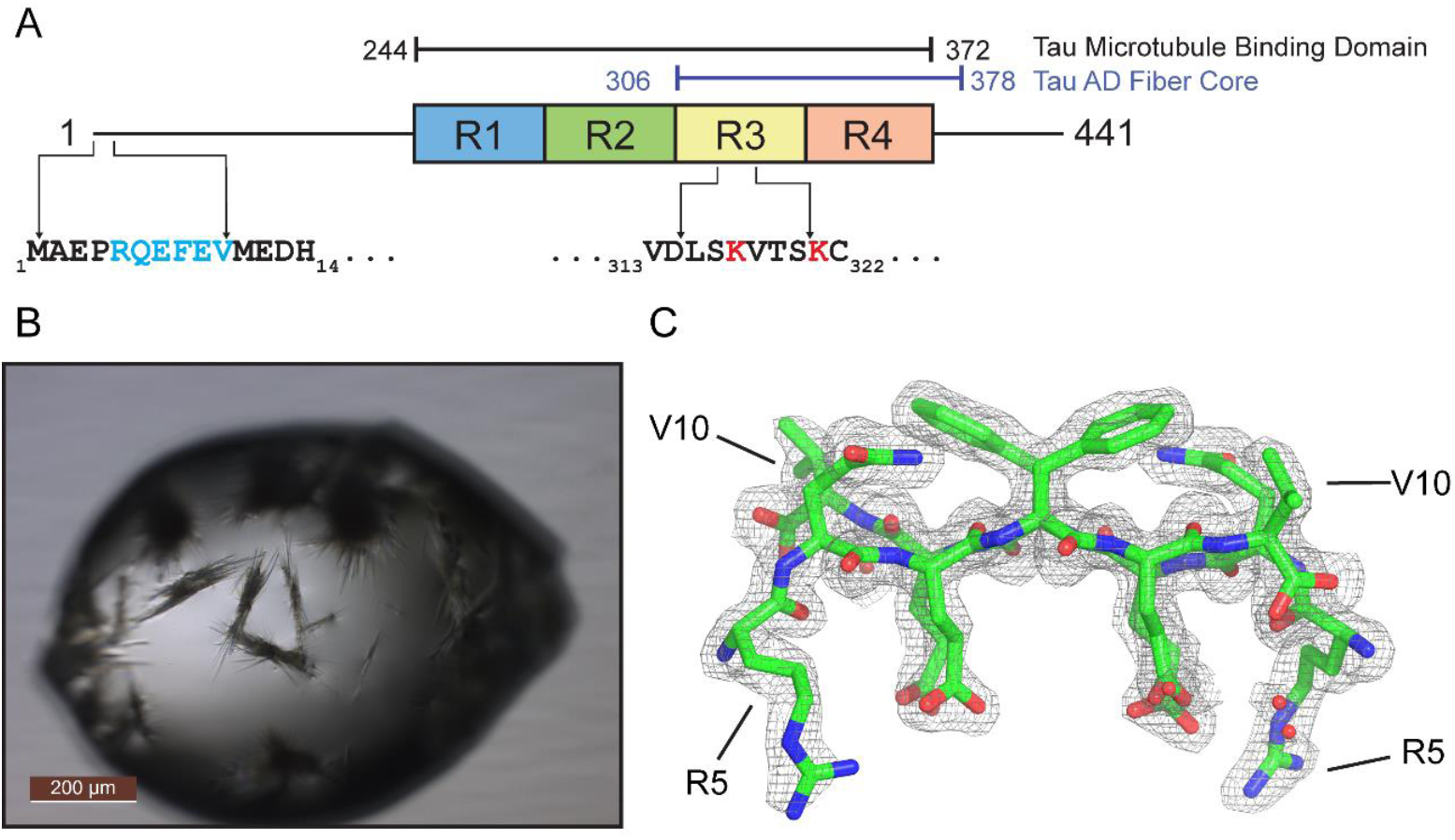
Crystal Structure of tau N-terminal segment _5_RQEFEV_10_. A) Schematic of tau primary structure. B) Crystals of _5_RQEFEV_10_ grown using the hanging drop method. C) Atomic model and electron density of _5_RQEFEV_10_ demonstrating the quality of fit. The view is down the fibril axis, showing two anti-parallel strands.

The crystal structure of _5_RQEFEV_10_ revealed a Class 5 homozipper where beta-strands assemble in antiparallel sheets and these sheets mate together in distinct face-to-face and back-to-back interfaces. Notably, the sheets are out-of-register and are related to each other by a 2_1_ “fibril axis” (20) (Figure 2 A, B). This combination of symmetry elements produces an ∼80° crossing angle between strands of one sheet and its mated sheet (Figure 2 A)(21). The alternating sequence of charged and hydrophobic/uncharged residues leads to wet and dry interfaces in the crystal structure.

**Figure 2:**
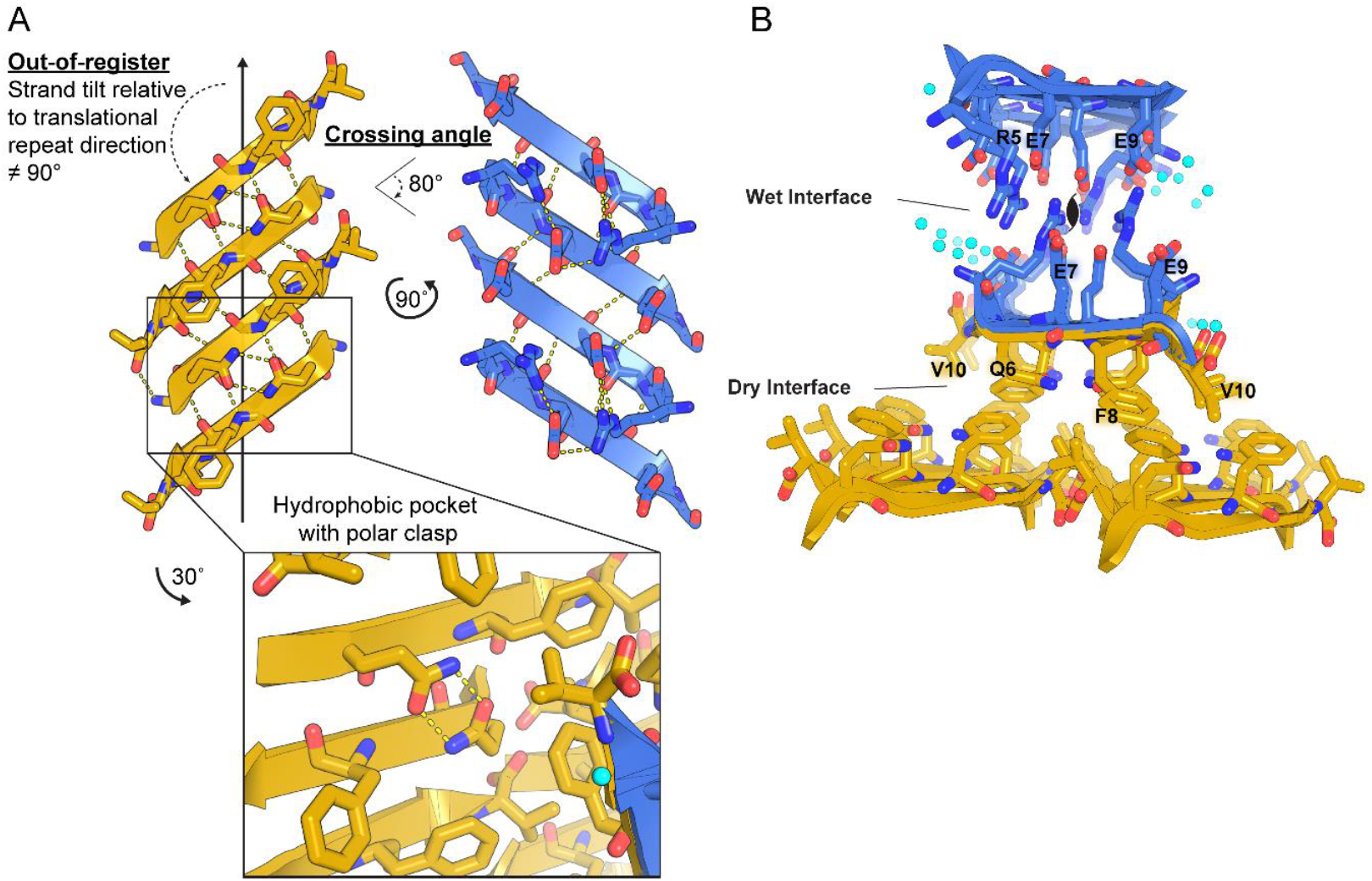
Crystal structure of _5_RQEFEV_10_ reveals a wet and a dry interface. A) _5_RQEFEV_10_ forms amyloid-like out-of-register protofilaments with wet and dry interfaces. Inset shows formation of a polar clasp with neighboring glutamines in the hydrophobic pocket of the dry interface. B) View down the fibril axis of _5_RQEFEV_10_ highlighting the interactions between residues within the wet and dry interfaces. Water molecules are shown by aqua spheres.

The wet interface features electrostatic interactions among polar, charged residues and water molecules. In particular, glutamates form an extensive hydrogen bond network with water molecules and arginines originating from the same sheet and from the opposing sheet (Figure 2 B). The dry interface features hydrophobic packing of phenylalanine, glutamine, and valine leading to the exclusion of water (Figure 2 A, B). Also, glutamine side chains clasp each other through a pair of hydrogen bonds, further stabilizing connections between neighboring strands in a sheet (Figure 2 A). This interaction is similar to the polar clasp described by Gallagher-Jones, *et al.*, with the distinction that glutamines in that study originated within the same strand(22). Similar to that polar clasp, neighboring aromatic residues restrict the glutamines to a conformation in which they bond to each other within a hydrophobic pocket (Figure 2 A). As stated by Gallagher-Jones, *et al.* the shielding of glutamines by neighboring aromatic residues may be essential for the formation of this polar clasp.

The crystal structure of _5_RQEFEV_10_ can account for the low resolution density found in the cryo-EM reconstructions of Alzheimer’s Disease (AD) tau filaments near residues K317 and K321, much as suggested by Fitzpatrick, *et al* (12). The positioning of _5_RQEFEV_10_ near these residues in the tau filament conformation is supported by the binding of the MC-1 and Alz50 antibodies to a discontinuous epitope consisting of both _7_EFE_9_ and _313_VDLSKVTSKC_322_(17).

In order to examine the potential interaction of the N-terminal _7_EFE_9_ segment with the AD fibril core, we first computationally docked the _6_QEFEV_10_ segment seen in the crystal structure into the low-resolution density shown to be adjacent to residues K317 and K321 in the AD Paired Helical Filament (PHF) (Figure 3 A-B)(12). In this model, the wet interface glutamates found in the crystal structure form electrostatic interactions with the exposed lysines in the PHF fibril, while the dry interface faces away from the PHF surface (Figure 3 A, B). Notably, we omitted Arg5 in this model due to steric clashes with Leu315 on the PHF. We speculate that Arg5 would have to adopt a different conformation in the fibril structure than in the crystal structure in order to maintain the interaction of Glu7 and Glu9 with Lys317 and Lys321.

**Figure 3:**
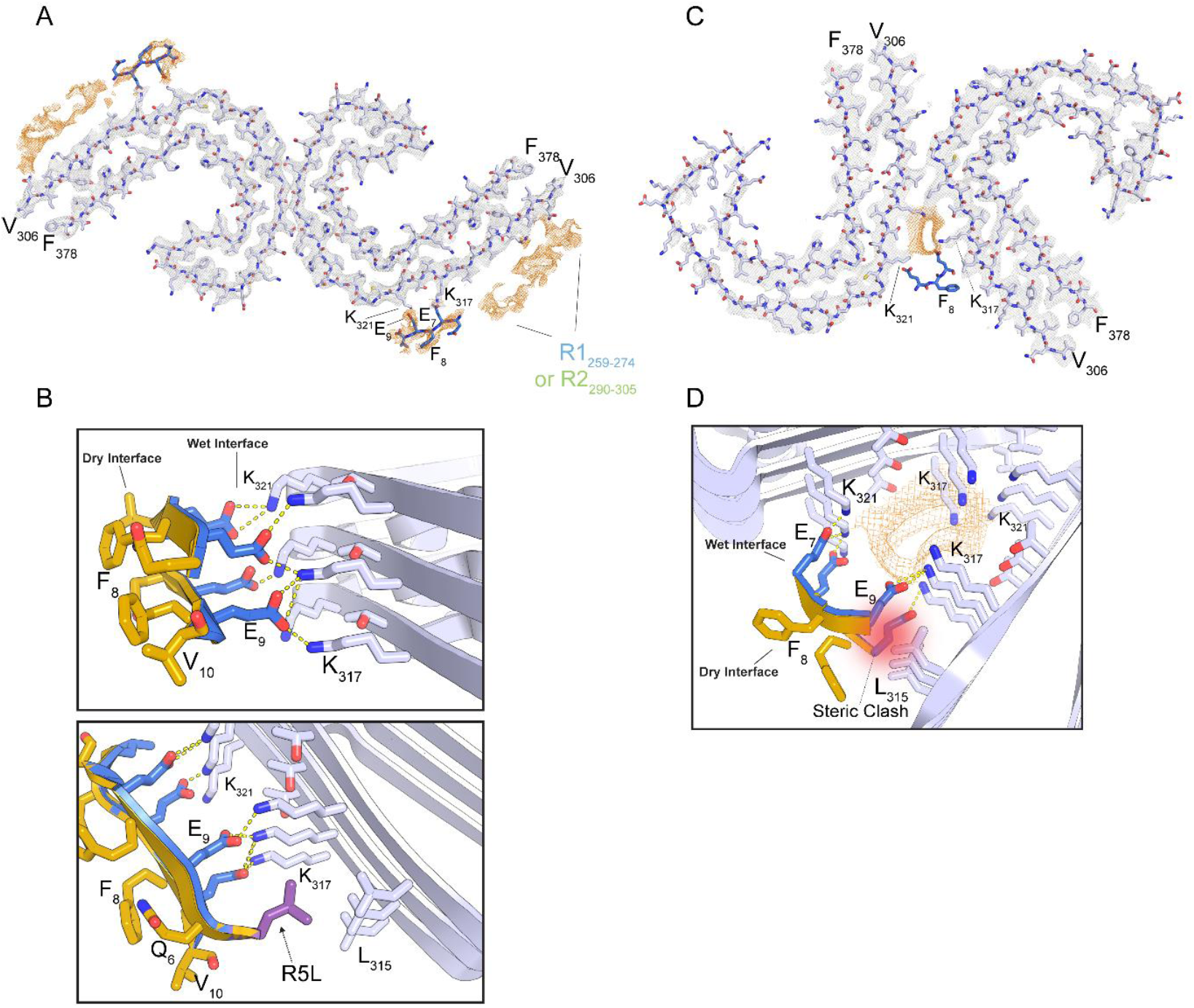
Speculative model for _5_RQEFEV_10_ interaction with Alzheimer’s Disease paired helical and straight filaments fibril cores. A) Atomic model of Alzheimer’s disease paired helical filaments (PHF) (5o3l.pdb) shown with electron density of modeled (grey) and un-modeled (orange) regions(12). _5_RQEFEV_10_ is docked into un-modeled density flanking the solvent-exposed K317 and K321 residues of the PHF. B) Detail (top) highlighting the interaction of the glutamates in the wet interface with K317 and K321 of the PHF. Detail (bottom) demonstrating the possible interaction of the R5L mutation with L315 of the PHF. C) Overview of potential interaction of _7_EFE_9_ with straight filaments (SF) (5o3t.pdb) at the inter-protofilament interface. D) Detail of the potential hydrogen bonding of wet interface glutamates with K317 and K321 and potential steric clash with L315 of the SF.

To examine further the relevance of the _7_EFE_9_ and _313_VDLSKVTSKC_322_ interaction in tau fibrils, we searched the literature for other evidence that implicates the N-terminus of tau in fibril formation. Poorkaj, P. *et al.* described a missense mutation found in a Progressive Supranuclear Palsy (PSP) patient that changes R5 to a leucine(23). In addition, it has been shown that deletion of residues 2-18 produces less aggregated tau than the wild-type sequence whereas the inclusion of the R5L mutation increases the amount of aggregated tau in the presence of arachidonic acid(24). This is consistent with our model of _7_EFE_9_ binding to _313_VDLSKVTSKC_322_ in the AD PHF; in that the deletion of residues 2-18 would abrogate the interaction of _7_EFE_9_ with _313_VDLSKVTSKC_322_. In addition, in our model the R5L mutation would result in a more stable interaction with Leu315 as discussed below.

To analyze if the R5L mutation might affect the binding of _5_RQEFEV_10_ to the _313_VDLSKVTSKC_322_ region in the AD filaments, we modeled the putative interaction of the sequence _5_LQEFEV_10_ with the AD PHF. To accomplish this, we mutated the R5 that was omitted in the wild-type model due a potential steric clash with L315 on the PHF, to a rotamer of leucine that would maximize its buried surface area and shape complementarity to L315 on the PHF (Figure 3 B). The model demonstrates that the mutation R5L would result in a more favorable interaction with the PHF than the native sequence, providing an explanation for R5L’s ability to increase tau aggregation.

Our attempts to dock the _5_RQEFEV_10_ crystal structure into the _313_VDLSKVTSKC_322_ region on the cryoEM structure of the straight filaments (SFs) were hindered due to the tight packing protofilaments that occurs in this region. By truncating the residues present in the crystal structure to only _7_EFE_9_ it is possible to place these residues within hydrogen bonding distance of K317 on one protofilament and K321 on the other protofilament. This results in a binding site comprised of residues from two different tau monomers, as opposed to a binding site comprised of only one monomer as in the PHF (Figure 3D). However, this two-tau monomer model of _7_EFE_9_ bound to the SF would result in steric clashes if any other residues were added to the _7_EFE_9_ sequence (Figure 3D), particularly with L315, making it harder to assess whether there is enough space in the SF inter-protofilament interface for the N-terminal _7_EFE_9_ sequence. Likewise, it was difficult to examine the effect of the R5L mutation on this interaction due to the resulting steric clashes.

## Discussion

The initial proposal that _7_EFE_9_ interacts with _313_VDLSKVTSKC_322_ came from biochemical studies in which Jicha, *et al.* confirmed that two antibodies, MC-1 and Alz50, most likely bind a single epitope of tau comprised of discontinuous segments _7_EFE_9_ and _313_VDLSKVTSKC_322_. The idea of a single epitope comprising these two distal sequences was supported by antibody binding assays using a series of tau constructs containing truncations or mutations in these regions(19). Tau constructs missing either _7_EFE_9_ or _313_VDLSKVTSKC_322_ did not exhibit antibody binding, demonstrating that both sequences need to be present for antibody reactivity. In addition, a series of mutations to the _7_EFE_9_ segment (Glu7,9 -> Ala7,9; Phe8 -> Ser8) abrogated antibody binding to tau. Importantly, Jicha, *et al* showed that tau constructs missing _7_EFE_9_ or _313_VDLSKVTSKC_322_ could not be mixed in solution to recover the MC-1/Alz50 epitope, indicating that this epitope is formed intramolecularly.

In an attempt to examine which sequences might interact with the primary nucleating sequences of tau _275_VQIINK_280_ and _306_VQIVYK_311_, Moore *et al.* tested the ability of different tau sequences to accelerate and increase _275_VQIINK_280_ and _306_VQIVYK_311_ aggregation(19). Through these experiments, the authors predicted the heterozipper interaction formed between _306_VQIVYK_311_ and _375_KLTFR_379_. This predicted interaction was later confirmed by the AD tau filament structure(12). In addition, Moore, *et al* showed that _5_RQEFEV_10_ can form fibrils *in vitro*(19), although it did not affect the aggregation of either _275_VQIINK_280_ or _306_VQIVYK_311_. This supports the idea that _5_RQEFEV_10_ instead interacts with _313_VDLSKVTSKC_322_ in a different region of the fibril core. Further experiments similar to those performed by Moore, *et al*, including aggregation kinetics and circular dichroism of the individual peptides and a mixture of both peptides, could help strengthen evidence for the interaction of _5_RQEFEV_10_ and _313_VDLSKVTSKC_322_ in the fibril state.

The results obtained by Jicha, *et al* and Moore, *et al* are consistent with the model proposed here where _5_RQEFEV_10_ occupies the un-modeled density that flanks residues K317 and K321 in the Fitzpatrick, *et al.* PHF cryo-em reconstruction(12). In particular, the abrogation of antibody binding by Glu7,9 -> Ala7,9 mutations performed by Jicha, *et al* can be explained by the disruption of the charge-charge interaction of glutamate and lysine residues in the proposed model (Figure 3 B)(17). The loss of this interaction would most likely greatly reduce the affinity of _7_EFE_9_ for the _313_VDLSKVTSKC_322_ segment, leading to the loss of the MC-1 and Alz50 epitope. The loss of antibody binding from the Phe8 -> Ser8 can be explained in the proposed model given that Phe8 is facing away from the fibril, allowing it to remain exposed for antibody binding. Therefore, mutation of Phe8 may not prevent the far N-terminal segment from binding to the exposed lysines on the fibril core, but may still eliminate antibody reactivity. This suggests that the _7_EFE_9_ segment needs to be not only in a stacked conformation bound to K317 and K321 on the fibril core, but also needs F8 to be facing away from the fibril core and presented for antibody binding. A loss of either of these conditions would result in a loss of MC-1 reactivity.

The model of _7_EFE_9_ interacting with K317 and K321 in the SF (Figure 3 C, D) suggests that either the _7_EFE_9_ sequence binds in a different manner to the _313_VDLSKVTSKC_322_ region on the SF or that the un-modeled density present in the Fitzpatrick, *et al* SF reconstruction does not result from the binding of the _7_EFE_9_ motif, but perhaps some other anion. Tau AD filament structures from 3 additional cases seem to recapitulate the extra density seen at the SF inter-protofilament interface(13). This indicates that this density may be a common feature of the SF fold and necessary to interact with the four lysines resulting from K317 and K321 of each protofilament coming together at the SF inter-protofilament interface.

It is worth noting that the cryo-EM structures of AD tau fibrils display parallel, in-register beta-strands, whereas the RQEFEV crystal structure forms out-of-register, antiparallel beta-sheets. Because residues N-terminal to Val306 are not resolved in the cryo-EM structure, we cannot determine whether _5_RQEFEV_10_ stacks into parallel or antiparallel sheets in the fibril. Our model used two strands of _5_RQEFEV_10_ stacked in an anti-parallel beta-sheet as seen in the crystal structure. Although different from the crystal structure, parallel, in-register beta-sheets of _5_RQEFEV_10_ would still form a wet and dry interface due to the alternating sequence of hydrophilic, charged residues and uncharged, mostly hydrophobic residues. Therefore, a parallel, in-register conformation of _5_RQEFEV_10_ would still allow Glu7 and Glu9 to form electrostatic interactions with Lys317 and Lys321 in a manner similar to the model proposed in Figure 3 A-B.

Recently, a new polymorph of tau from the brain of a Pick’s Disease case has been visualized by cryo-EM(14). This structure adopts a drastically different fold from the AD filaments; however, the Pick’s Disease filaments are still MC-1 reactive, indicating the preservation of the _7_EFE_9_ and _313_VDLSKVTSKC_322_ epitope(14). In this structure, K317 and K321 are exposed to the solvent in a beta-sheet conformation, which would allow the N-terminal _7_EFE_9_ segment to bind K317 and K321 through electrostatic interactions between the glutamates and lysines similar to the AD PHF model (Figure 3 A, B). This electrostatic interaction would preserve the MC-1 epitope and provide an explanation for why MC-1 recognizes both tau fibril polymorphs.

In addition, the potential strengthening of the N-terminal interaction with the fibril core through the R5L mutation and its discovery in a PSP patient, suggests that this interaction may also occur in the PSP tau fibril. Although there is evidence that so-called 4R tauopathies, where the dominant species found in aggregated tau are the 4R isoforms, PSP and Corticobasal Degeneration (CBD) form different tau polymorphs, their structures have not yet been determined(25). However, as long as the _313_VDLSKVTSKC_322_ region adopts a beta-sheet like fold, and K317 and K321 remain solvent-exposed, the long-range charge-charge interaction with _7_EFE_9_ could be preserved. In short, there may be a common interaction among the disparate folds of tau polymorphs.

In the past, our lab has developed inhibitors of tau aggregation by structure-based drug design(11,16). This requires detailed structural knowledge of a site of the tau protein in the aggregated state obtained by X-ray crystallography or MicroED. These inhibitors target segments of the tau protein in the microtubule binding region that is thought to participate in the fibril core of all tau filaments. However, given the structural evidence thus far that the microtubule binding region can adopt different folds in different diseases, it is likely that a spectrum of inhibitors will be necessary to most effectively block aggregation or spreading of specific tau polymorphs. Immuno-labeling with MC-1 seems to indicate that the N-terminal interaction with the fibril core modeled here is preserved in both AD and Pick’s Disease tau filaments. Therefore, an inhibitor targeted towards this interaction may be general to all tau filaments, providing another target for treating tauopathies.

## Methods

### Crystallization and Data Collection

Synthetic peptide RQEFEV was ordered from GenScript. RQEFEV was crystallized using the hanging drop method with a 2:1 mixture of 60 mg/mL RQEFEV and 0.2 M Ammonium Citrate Dibasic, 30% PEG 3350. Diffraction data was collected at APS Beamline 24-ID-E using an Eiger detector. **Data Processing and Structure**

### Data Processing and Structure Determination

Diffraction data were indexed and integrated using XDS and scaled using XSCALE(26). Molecular replacement was performed using Phaser and an idealized beta-strand as a molecular replacement probe(27). Model-building and manual real-space refinement was performed in COOT(28). Automated reciprocal-space and real-space refinement was performed using Refmac and Phenix(29, 30).

### Modeling

Modeling was performed in COOT using the RQEFEV crystal structure and the cryo-em structures for the AD PHF (5o3l.pdb) and SF (5o3t.pdb) downloaded from the PDB. Cryo-em maps for the PHF (EMD-3741) and SF (EMD-3743) were also used for modeling and generating figures. All figures were made in Pymol (Schrodinger).

**Table 1.** Data collection and refinement statistics.

|  | RQEFEV (PDB ID: 6N4P) |
| --- | --- |
| Wavelength | 0.9792 |
| Resolution range | 16.08 - 1.851 (1.918 - 1.851) |
| Space group | P2<sub>1</sub> |
| Unit cell | 16.59 11.45 25.42 90 104.236 90 |
| Total reflections | 2416 (226) |
| Unique reflections | 842 (80) |
| Multiplicity | 2.9 (2.8) |
| Completeness (%) | 97.2 (96.4) |
| Mean I/sigma(I) | 4.2 (1.6) |
| Wilson B-factor | 14.8 |
| R-merge | 0.16 (0.60) |
| R-meas | 0.20 (0.72) |
| R-pim | 0.11 (0.40) |
| CC1/2 | 0.97 (0.83) |
| CC* | 0.99 (0.95) |
| Reflections used in refinement | 838 (80) |
| Reflections used for R-free | 85 (8) |
| R-work | 0.19 (0.30) |
| R-free | 0.27 (0.47) |
| CC(work) | 0.96 (0.81) |
| CC(free) | 0.91 (0.83) |
| Number of non-hydrogen atoms | 118 |
| macromolecules | 114 |
| solvent | 4 |
| Protein residues | 12 |
| RMS(bonds) | 0.013 |
| RMS(angles) | 1.48 |
| Ramachandran favored (%) | 100.00 |
| Ramachandran allowed (%) | 0.00 |
| Ramachandran outliers (%) | 0.00 |
| Rotamer outliers (%) | 0.00 |
| Clashscore | 4.55 |
| Average B-factor | 21.9 |
| macromolecules | 21.5 |
| solvent | 32.5 |

Statistics for the highest-resolution shell are shown in parentheses

## Acknowledgments

We thank Michael Sawaya for discussion and Michael Collazo at UCLA-DOE Macromolecular Crystallization Core Technology Center for crystallization support. We thank NIH AG 054022. This work is based upon research conducted at the Northeastern Collaborative Access Team beamlines, which is funded by the National Institute of General Medical Sciences from the National Institutes of Health (P41 GM103403). The Eiger 16M detector on 24-ID-E beam line is funded by a NIHORIP HEI grant (S10OD021527). This research used resources of the Advanced Photon Source, a U.S. Department of Energy (DOE) Office of Science User Facility operated for the DOE Office of Science by Argonne National Laboratory under Contract No. DE-AC02-06CH11357. DRB is funded by the National Science Foundation Graduate Research Fellowship Program.

## References

1. Kadavath H, Hofele RV, Biernat J, Kumar S, Tepper K, Urlaub H, et al. Tau stabilizes microtubules by binding at the interface between tubulin heterodimers. Proc Natl Acad Sci U S A. 2015 Jun 16;112(24):7501–6.

2. Kellogg EH, Hejab NMA, Poepsel S, Downing KH, DiMaio F, Nogales E. Near-atomic model of microtubule-tau interactions. Science. 2018 May 10;eaat1780.

3. Miyata Y, Koren J, Kiray J, Dickey CA, Gestwicki JE. Molecular chaperones and regulation of tau quality control: strategies for drug discovery in tauopathies. Future Med Chem. 2011 Sep;3(12):1523–37.

4. Goedert M, Wischik CM, Crowther RA, Walker JE, Klug A. Cloning and sequencing of the cDNA encoding a core protein of the paired helical filament of Alzheimerdisease: identification as the microtubule-associated protein tau. Proc Natl Acad Sci. 1988 Jun 1;85(11):4051–5.

5. Wischik CM, Novak M, Thøgersen HC, Edwards PC, Runswick MJ, Jakes R, et al. Isolation of a fragment of tau derived from the core of the paired helical filament of Alzheimer disease. Proc Natl Acad Sci U S A. 1988 Jun;85(12):4506–10.

6. Wischik CM, Novak M, Edwards PC, Klug A, Tichelaar W, Crowther RA. Structural characterization of the core of the paired helical filament of Alzheimer disease. Proc Natl Acad Sci U S A. 1988 Jul;85(13):4884–8.

7. Narayanan RL, Dürr UHN, Bibow S, Biernat J, Mandelkow E, Zweckstetter M. Automatic Assignment of the Intrinsically Disordered Protein Tau with 441-Residues. J Am Chem Soc. 2010 Sep 1;132(34):11906–7.

8. Jeganathan S, von Bergen M, Brutlach H, Steinhoff H-J, Mandelkow E. Global Hairpin Folding of Tau in Solution. Biochemistry. 2006 Feb 1;45(7):2283–93.

9. Mirbaha H, Chen D, Morazova OA, Ruff KM, Sharma AM, Liu X, et al. Inert and seed-competent tau monomers suggest structural origins of aggregation. Akhmanova A, editor. eLife. 2018 Jul 10;7:e36584.

10. Sawaya MR, Sambashivan S, Nelson R, Ivanova MI, Sievers SA, Apostol MI, et al. Atomic structures of amyloid cross-β spines reveal varied steric zippers. Nature. 2007 May;447(7143):453–7.

11. Seidler PM, Boyer DR, Rodriguez JA, Sawaya MR, Cascio D, Murray K, et al. Structure-based inhibitors of tau aggregation. Nat Chem. 2018 Feb;10(2):170–6.

12. Fitzpatrick AWP, Falcon B, He S, Murzin AG, Murshudov G, Garringer HJ, et al. Cryo-EM structures of tau filaments from Alzheimer’s disease. Nature. 2017 Jul 13;547(7662):185–90.

13. Falcon B, Zhang W, Schweighauser M, Murzin AG, Vidal R, Garringer HJ, et al. Tau filaments from multiple cases of sporadic and inherited Alzheimer’s disease adopt a common fold. Acta Neuropathol (Berl) [Internet]. 2018 Oct 1 [cited 2018 Oct 25]; Available from: https://doi.org/10.1007/s00401-018-1914-z

14. Falcon B, Zhang W, Murzin AG, Murshudov G, Garringer HJ, Vidal R, et al. Structures of filaments from Pick’s disease reveal a novel tau protein fold. Nature. 2018 Sep;561(7721):137.

15. Bergen M von, Friedhoff P, Biernat J, Heberle J, Mandelkow E-M, Mandelkow E. Assembly of τ protein into Alzheimer paired helical filaments depends on a local sequence motif (306VQIVYK311) forming β structure. Proc Natl Acad Sci. 2000 May 9;97(10):5129–34.

16. Sievers SA, Karanicolas J, Chang HW, Zhao A, Jiang L, Zirafi O, et al. Structure-based design of non-natural amino-acid inhibitors of amyloid fibril formation. Nature. 2011 Jun 15;475(7354):96–100.

17. Jicha GA, Bowser R, Kazam IG, Davies P. Alz-50 and MC-1, a new monoclonal antibody raised to paired helical filaments, recognize conformational epitopes on recombinant tau. J Neurosci Res. 1997 Apr 15;48(2):128–32.

18. Thompson MJ, Sievers SA, Karanicolas J, Ivanova MI, Baker D, Eisenberg D. The 3D profile method for identifying fibril-forming segments of proteins. Proc Natl Acad Sci. 2006 Mar 14;103(11):4074–8.

19. Moore CL, Huang MH, Robbennolt SA, Voss KR, Combs B, Gamblin TC, et al. Secondary nucleating sequences affect kinetics and thermodynamics of tau aggregation. Biochemistry. 2011 Dec 20;50(50):10876–86.

20. Eisenberg DS, Sawaya MR. Structural Studies of Amyloid Proteins at the Molecular Level. Annu Rev Biochem. 2017 20;86:69–95.

21. Liu C, Zhao M, Jiang L, Cheng P-N, Park J, Sawaya MR, et al. Out-of-register β-sheets suggest a pathway to toxic amyloid aggregates. Proc Natl Acad Sci. 2012 Dec 18;109(51):20913–8.

22. Poorkaj P, Muma NA, Zhukareva V, Cochran EJ, Shannon KM, Hurtig H, et al. An R5L τ mutation in a subject with a progressive supranuclear palsy phenotype. Ann Neurol. 2002 Oct 1;52(4):511–6.

23. Gamblin TC, Berry RW, Binder LI. Tau polymerization: role of the amino terminus. Biochemistry. 2003 Feb 25;42(7):2252–7.

24. Gibbons GS, Banks RA, Kim B, Changolkar L, Riddle DM, Leight SN, et al. Detection of Alzheimer Disease (AD)-Specific Tau Pathology in AD and NonAD Tauopathies by Immunohistochemistry With Novel Conformation-Selective Tau Antibodies. J Neuropathol Exp Neurol. 2018 Mar 1;77(3):216–28.

25. Kabsch W. XDS. Acta Crystallogr D Biol Crystallogr. 2010 Feb 1;66(Pt 2):125–32.

26. McCoy AJ, Grosse-Kunstleve RW, Adams PD, Winn MD, Storoni LC, Read RJ. Phaser crystallographic software. J Appl Crystallogr. 2007 Aug 1;40(Pt 4):658–74.

27. Emsley P, Lohkamp B, Scott WG, Cowtan K. Features and development of Coot. Acta Crystallogr D Biol Crystallogr. 2010 Apr;66(Pt 4):486–501.

28. Murshudov GN, Vagin AA, Dodson EJ. Refinement of macromolecular structures by the maximum-likelihood method. Acta Crystallogr D Biol Crystallogr. 1997 May 1;53(Pt 3):240–55.

29. Afonine PV, Grosse-Kunstleve RW, Echols N, Headd JJ, Moriarty NW, Mustyakimov M, et al. Towards automated crystallographic structure refinement with phenix.refine. Acta Crystallogr D Biol Crystallogr. 2012 Apr 1;68(4):352–67.

